# RetINaBox: A hands-on learning tool for experimental neuroscience

**DOI:** 10.1101/2025.09.11.673547

**Authors:** Brune Bettler, Flavia Arias Armas, Erica Cianfarano, Vanessa Bordonaro, Megan Liu, Matthew Loukine, Mingyu Wan, Aude Villemain, Blake Richards, Stuart Trenholm

## Abstract

An exciting aspect of neuroscience is developing and testing hypotheses via experimentation. However, due to logistical and financial hurdles, the experiment and discovery component of neuroscience is generally lacking in classroom and outreach settings. To address this issue, here we introduce RetINaBox: a low-cost open-source electronic visual system simulator that provides users with a hands-on tool to discover how the visual system builds feature detectors. RetINaBox features an LED array for generating visual stimuli and a photodiode array that acts as a mosaic of model photoreceptors. Custom software on a Raspberry Pi computer reads out responses from model photoreceptors and allows users to control the polarity and delay of the signal transfer from model photoreceptors to model retinal ganglion cells. Interactive lesson plans are provided, guiding users to discover different types of visual feature detectors—including ON/OFF, center-surround, orientation selective, and direction selective receptive fields—as well as their underlying circuit computations.

## Introduction

The manner in which the brain encodes sensory stimuli varies across brain areas and is often far from predictable^1–5^. Thus, to discover how sensory inputs are represented in the brain we need to directly record from individual neurons while providing sensory stimulation^5^. A challenge in neuroscience education and outreach is how to incorporate such experimental work into lesson plans, so that instead of solely learning lists of facts, students get hands-on experience that captures the excitement of discovery. Laboratory classes have long been used to address this challenge, but their scope is often limited by financial, infrastructural, technical, ethical, and training constraints. For example, *in vivo* single-unit recordings from animal brains during sensory stimulation are widely used to capture the neural code but are, for the most part, too difficult to recapitulate in pedagogical settings.

To this end, we developed RetINaBox (“retina in a box”) as a neuroscience focused educational and outreach tool that provides users with an interactive, hands-on system with which to discover how the visual system implements several important feature selective computations that were discovered through single-cell neurophysiological recordings^5^. RetINaBox consists of a low-cost computer, simple electronic components (LEDs and photodiodes), 3D printed parts, and custom-written open-source software. Through several lesson plans, RetINaBox exposes users to numerous computations in the visual system, including ON/OFF processing^6–8^, center-surround receptive fields^9,10^, orientation selectivity^3,11^ and direction selectivity^12,13^. RetINaBox also includes Discovery Mode, which recreates the experience of being an experimental visual neuroscientist: users load a preset neuron into RetINaBox but are not shown the wiring scheme of its inputs; users then need to test different visual stimuli until they discover the specific stimulus that activates the neuron; finally, users are tasked with discovering the circuit wiring scheme that underlies the neuron’s stimulus selectivity.

## Materials and Methods

RetINaBox comprises an electronic visual stimulator and retina combined with software to readout and control incoming visual signal in model visual circuits. A complete parts list for building RetINaBox is provided in the user manual, which is included in the supplementary materials and on Github. The user manual also includes a step-by-step guide for building RetINaBox. To ensure that RetINaBox is as cost-effective as possible, we built it around a Raspberry Pi 500, which features a Raspberry Pi 5 minicomputer integrated into a keyboard, with easy access to GPIO pins for sending signals to external devices and monitoring external signals. We provide custom-written open-source software for running RetINaBox. The user manual includes instructions for software installation and use. At the time of publication, the cost for all components, including a Raspberry Pi and a monitor, was < $350 USD. The build time (once users have all components and 3D printed parts in hand) is ~4 hours.

## Results

### Design of RetINaBox

We sought to develop a simplified model retina that could be paired with a computer to allow users to explore, via experimentation and discovery, how visual inputs detected by retinal photoreceptors are processed in downstream visual neurons to generate feature selective responses. To do so, we designed a simplified electronic retina, with a built-in visual stimulator, and connected it to a Raspberry Pi computer loaded with software for simulating neuronal circuit processing.

#### Simplified model retina

To build RetINaBox, we sacrificed some biological details, since our main goal was to introduce users to general principles without requiring them to first learn, one at a time, about specific implementations of these feature selective computations in specific cell types of the visual system. As such, and as outlined in more detail below and in **Figure 1A,B**, RetINaBox does not include model horizontal cells, bipolar cells, or amacrine cells.

**Figure 1.**
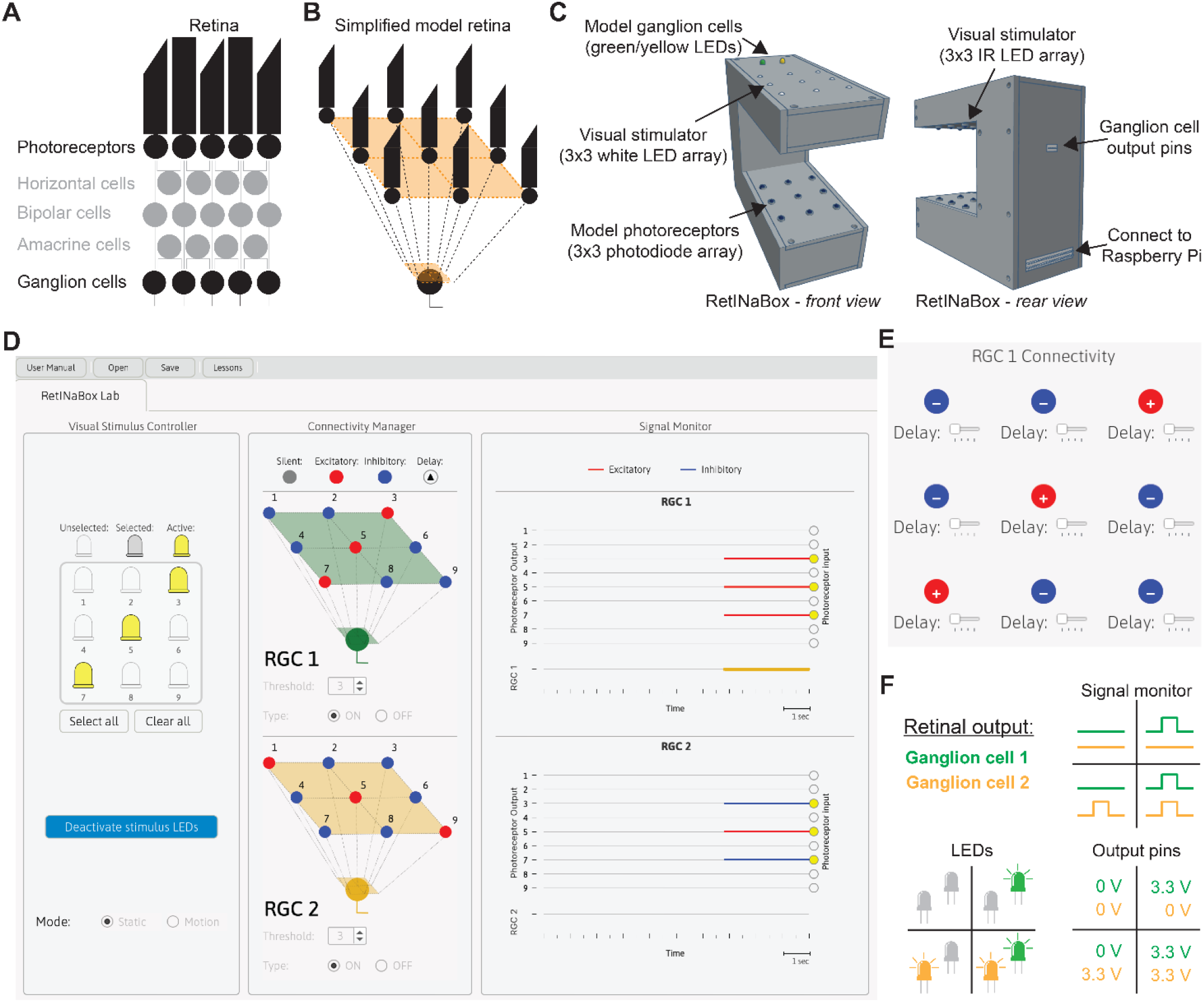
RetINaBox design and GUI overview. A schematic of retinal circuitry (**A**) and the simplified model circuitry of RetINaBox (**B**). **C**, An overview of RetINaBox, including the 3D printed case, the visual stimulator (3×3 LED array) and photoreceptor mosaic (3×3 photodiode array), along with the connectivity port between RetINaBox and the Raspberry Pi, and the output signals of the two model retinal ganglion cells (2 color LEDs on top of RetINaBox and 3.3V output pins on the back of RetINaBox). **D**, The RetINaBox GUI, showing the Stimulus Controller, Connectivity Manager, and Signal Monitor. **E**, The Connectivity Manager pop-up window, which allows users to set the connectivity (silent, +, or −) and delay (none, short, medium, or long) of the connection between each of the 9 model photoreceptors and each of the two retinal ganglion cells (RGCs). The Connectivity Manager allows users to set the RGC threshold (i.e. how many + inputs an RGC needs to receive to respond) and type (ON or OFF). **F**, Methods for monitoring RGC activity, either by watching the RGC responses in the GUI’s Signal Monitor, or by monitoring the yellow and green LEDs on the top of RetINaBox, or by connecting external electronic devices to the 3.3V output pins on the back of RetINaBox.

To provide a specific example of the pedagogical philosophy behind RetINaBox: instead of having six separate lessons outlining the exact biological details behind how center-surround is differentially implemented in 1) photoreceptors, 2) horizontal cells, 3) bipolar cells, 4) amacrine cells, 5) ganglion cells and 6) LGN neurons, RetINaBox simply includes a single lesson focused on the general concept of center-surround. As another example of our philosophy: while orientation selectivity and direction selectivity are implemented in various ways depending on cell type, brain area, and species^14,15^, with RetINaBox we have a single lesson for orientation selectivity and a single lesson for direction selectivity, where we introduce the general concepts.

Seeing starts in photoreceptors. In the vertebrate retina, photoreceptors form a single layer of cell bodies and are spatially organized as a lattice, or mosaic, next to one another, such that the light-receptive outer segments of neighbouring photoreceptors detect changes in light intensity in neighbouring regions of visual space^16–18^. To model photoreceptors, we used photodiodes, semiconductor diodes sensitive to changes in photon flux. To model a mosaic of photoreceptors, we built a 3 × 3 array of photodiodes (**Figure 1B,C**). This is the smallest photoreceptor array that can implement all the feature selective computations we sought to explore with RetINaBox: ON/OFF, center-surround, orientation selectivity and direction selectivity.

Next, while the vertebrate retina is a complex tissue comprised of multiple cell types located in specific anatomical layers^16^ (**Figure 1A**), here we designed a simplified model retina whereby the array of 9 model photoreceptors that can be connected to 2 model retinal ganglion cells (RGCs; **Figure 1B**). Users should note that this is a simplification—in the actual vertebrate retina photoreceptors do not directly connect to RGCs (**Figure 1A**). The contribution from other retinal neurons to visual computations are incorporated into the connectivity functions in RetINaBox that transform signals passing from photoreceptors to RGCs (as outlined in detail below). The photodiodes are powered by the Raspberry Pi, and the output of each photodiode is sampled by a different GPIO pin from the Raspberry Pi.

#### Visual stimulation

So that each photodiode can be independently activated, we aligned the photodiode array with a 3 × 3 LED array (**Figure 1C**), with each LED being independently controlled by a different GPIO pin from the Raspberry Pi. To ensure that photodiode activation is selectively modulated by the stimulation LEDs and not from variations in ambient light levels, we used infrared (IR) stimulation LEDs and IR-sensitive photodiodes. However, so that users can see the pattern and timing of LED stimulation on RetINaBox (the IR LEDs being invisible to the human eye), we added a second 3 × 3 array of visible (white light) LEDs pointing upward (i.e. away from the photodiodes; **Figure 1C**).

### Graphical user interface for ‘performing experiments’

To control LED stimulation, monitor model photoreceptor responses, connect model photoreceptors to model retinal ganglion cells, and monitor RGC output responses, we designed a graphical user interface (GUI) to control RetINaBox (**Figure 1D,E**). The GUI has 3 panels: a) Visual Stimulus Controller; b) Connectivity Manager; c) Signal Monitor (**Figure 1D**).

The Visual Stimulus Controller (**Figure 1D**) allows the user to independently control activation of each LED in the 3 × 3 array. Visual stimuli can be displayed in a static manner or can be made to move either leftward or rightward at 3 different speeds. As described in the lesson plan document, for initial testing of circuits in RetINaBox we recommend using the Visual Stimulus Controller to deliver specific patterns of light to the model photoreceptors. However, once the user is satisfied that they have generated appropriate feature selective circuits for a given activity, we recommend turning on all the stimulus LEDs and placing real-world stimuli between RetINaBox’s stimulus LEDs and photodiodes to test the robustness of the ganglion cell’s feature selectivity. Along these lines, in the user manual users are instructed to make shapes with modelling clay on a clear plastic sheet which serves as the Visual Stimulus Tool. Alternatively, users can use their hands or use pieces of paper/cardboard cut into various shapes to create visual stimuli.

The Connectivity Manager (**Figure 1D,E**) allows users to connect the output from each model photoreceptor to a model retinal ganglion cell. The GUI features 2 model retinal ganglion cells. By clicking on a model ganglion cell (**Figure 1D**), a pop-up allows users to modify the connectivity of each photoreceptor to that ganglion cell (**Figure 1E**). Each model ganglion cell can receive input from up to all 9 model photoreceptors. For each connection, the polarity (silent, +, or −; which corresponds to the photoreceptor sending either a 0, +1, or −1, respectively, to the RGC) and time delay (none, short, medium, long) can be adjusted. The response threshold for each ganglion cell can be set from between 1 to 9, indicating how many positive photoreceptor inputs, calculated in real-time by pooling all its inputs, it needs to receive to respond. Additionally, ganglion cells can be set as either ON or OFF type (**Figure 1D**), modeling ON/OFF retinal processing^16^. ON ganglion cells only receive signals from photoreceptors that are currently being stimulated by light, whereas OFF ganglion cells only receive signals from photoreceptors that are not currently being stimulated by light. Ganglion cell output is binary: on or off.

The Signal Monitor (**Figure 1D**) plots when each model photoreceptor (i.e. photodiode) is activated by the visual stimulus, the polarity of the signal transfer from each photoreceptor to each model ganglion cell, and each model ganglion cell’s output response (**Figure 1D,F**). In addition, the output signals of the 2 model retinal ganglion cells are indicated by 2 colored LEDs (RGC1, green; RGC2, yellow) on the top of RetINaBox (**Figure 1C,F**). Furthermore, the output of the ganglion cells can be used to drive external electronic components via output pins on the rear of RetINaBox that send out 3.3 V signals when a given model retinal ganglion cell is activated (**Figure 1C,F**).

### Lessons

Once users have built RetINaBox and installed the software on the Raspberry Pi (details are covered in the user manual), users are guided through 4 lesson plans. The first 3 lessons guide users through the circuit mechanisms underlying canonical visual system computations: ON/OFF and center-surround (lesson 1), orientation selectivity (lesson 2), and direction selectivity (lesson 3). Lesson 4 is focused on discovery, with users tasked with discovering the preferred visual stimulus of mystery RetINaBox ganglion cells and then discovering the circuit connectivity underlying each mystery cell’s feature selective tuning preference.

#### Lesson 1 – Explore ON/OFF processing and crack a code with center-surround receptive fields

Following experiments that found light-evoked spiking in the optic nerve^19^, retinal ganglion cells were found to have spatially-localized receptive fields (i.e. they only ‘see’ within a small part of the visual scene), with some cells responding only to increases in luminance over their receptive field (RF), some only responding to decreases in luminance, and some responding to both increases and decreases in luminance^6^. Such ON vs. OFF responses were subsequently found to arise at the level of bipolar cells, with ‘sign-conserving’ OFF bipolar cells possessing ionotropic glutamate receptors in their dendrites, and ‘sign-inverting’ ON bipolar cells possessing metabotropic glutamate receptors in their dendrites^16,20^. Such ON/OFF processing was found to play an important role in visual perception, helping with contrast sensitivity and enabling robust detection of both increases and decreases in luminance^21^.

It was also discovered that RGCs care about the pattern of visual stimuli falling within their receptive fields, due to a center-surround RF organization^5,9,10,16^. For example, it was found that for an ON-center retinal ganglion cell, increasing luminance with a spot of light located directly above the cell— usually corresponding to the location of its cell body and most of its dendritic tree—maximally activated the cell. However, if the size of the visual stimulus was increased beyond the receptive field center, the cell’s response decreased due to activation of an inhibitory surround. The same was found for OFF-center retinal ganglion cells, whose optimal stimulus is a luminance darkening over its receptive field center. While first described in RGCs, surround inhibition was subsequently described at each level of retinal processing—in photoreceptors^22^, horizontal cells^23^, bipolar cells^24^, and amacrine cells^25^—via inhibitory lateral connections, which in the vertebrate retina can arise from horizontal cells and amacrine cells^16^. Center-surround receptive fields mean that RGCs are optimized to respond to local luminance contrast^5^—meaning that most visual neurons are not strongly activated by homogenous, spatially redundant scenes—with the RF size relating to the optimal spatial frequency of luminance contrast that will activate a cell. It should be noted that, since for purpose of simplicity RetINaBox independently models ON and OFF systems, RetINaBox cannot model cross-over inhibition, in which ON and OFF retinal channels cross-inhibit one another, which can also modulate center-surround processing^27^.

Lesson 1 of RetINaBox asks users to consider the ecological purposes of ON/OFF and center-surround receptive fields, before asking users to build ganglion cells with ON vs. OFF response properties and center-surround receptive field organization. The first goal is to connect the photoreceptors to RGC1 so that it will respond when a single model photoreceptor is activated, but not when that photoreceptor is activated at the same time as other photoreceptors in the array (**Figure 2A-D**; e.g. generate an ON-center ganglion cell that is selective to a small spot of light centered on the middle of the photoreceptor array). The next goal is to generate a second ganglion cell with the same feature selectivity but located in a different part of the visual field (i.e. users will have two ON-center ganglion cells with RF centers in different locations). By generating 2 RGCs with receptive fields centered in different parts of the visual field, RetINaBox elucidates the point that an individual photoreceptor can contribute to distinct parts of different downstream neurons’ receptive fields (in this example the photoreceptor that is responsible for one RGC’s RF center contributes to the other RGC’s RF surround). The next goal involves switching one of the RGCs from ON-center to OFF-center. This will help users learn about how the visual system differentially processes increases vs. decreases in luminance (**Figure 2A-D**). Finally, users are tasked with generating two ON-center ganglion cells with the same receptive field center location, but with preferences for spots of different sizes (**Figure 3A-D**). This will show users how receptive field size controls the spatial frequency of luminance contrast that activates each RGC. If users need assistance solving these problems, they can consult the lesson plan document or check out example Connectivity Manager solutions which can be loaded as presets from the “Lessons” tab in the menu bar. In addition, the GUI allows users to save their solutions and access them later.

**Figure 2.**
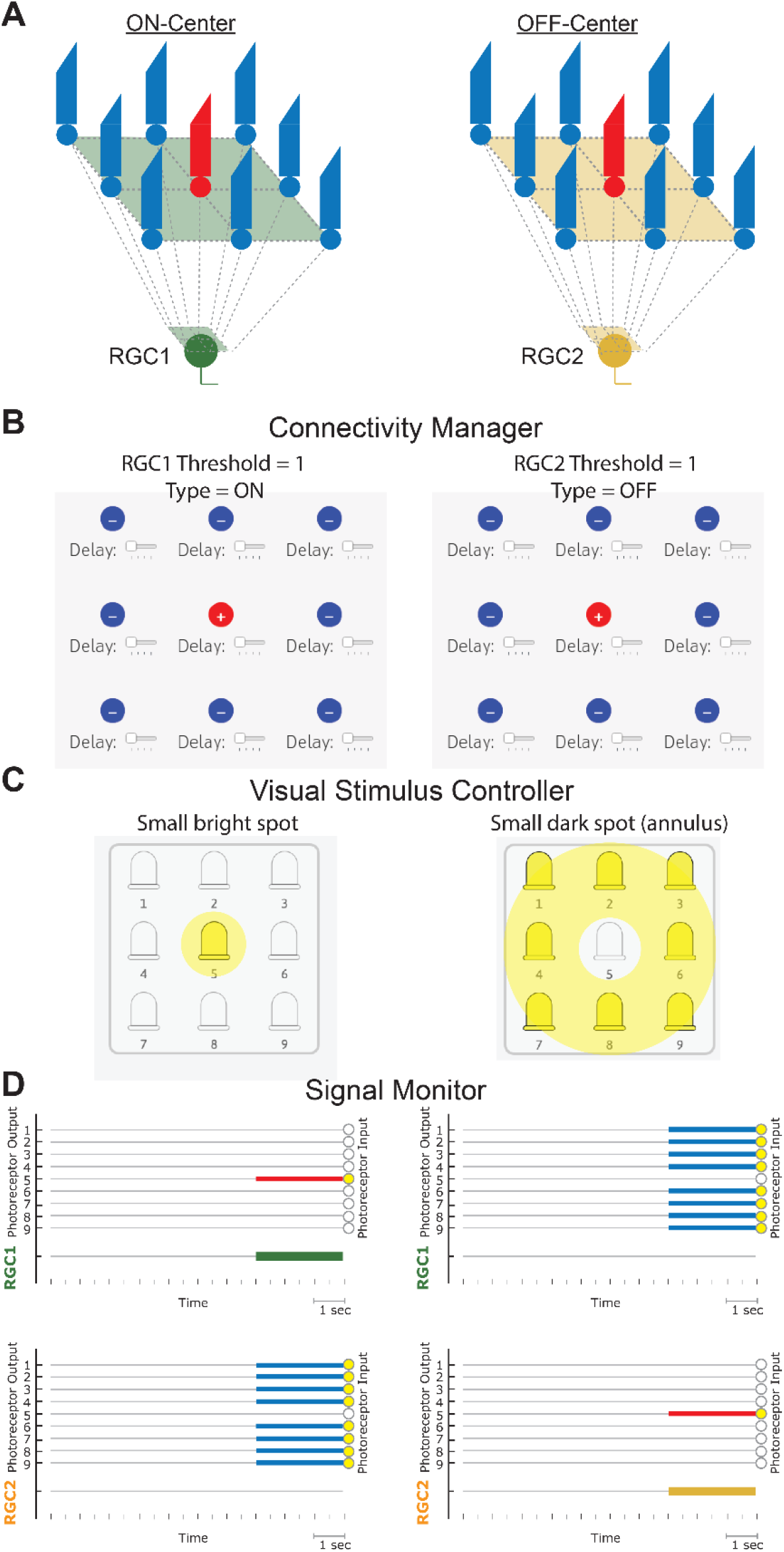
Lesson 1 – ON vs. OFF processing. **A**, Example circuits for generating RGCs with center-surround ON (*left*) and OFF (*right*) RFs. Example settings for the Connectivity Manager (**B**) and the Visual Stimulus Controller (**C**). **D**, Example Signal Monitor output for the settings outlined above, with the stimulus LEDs turned on for 3 s (small light spot, *left*; small dark spot, *right*).

**Figure 3.**
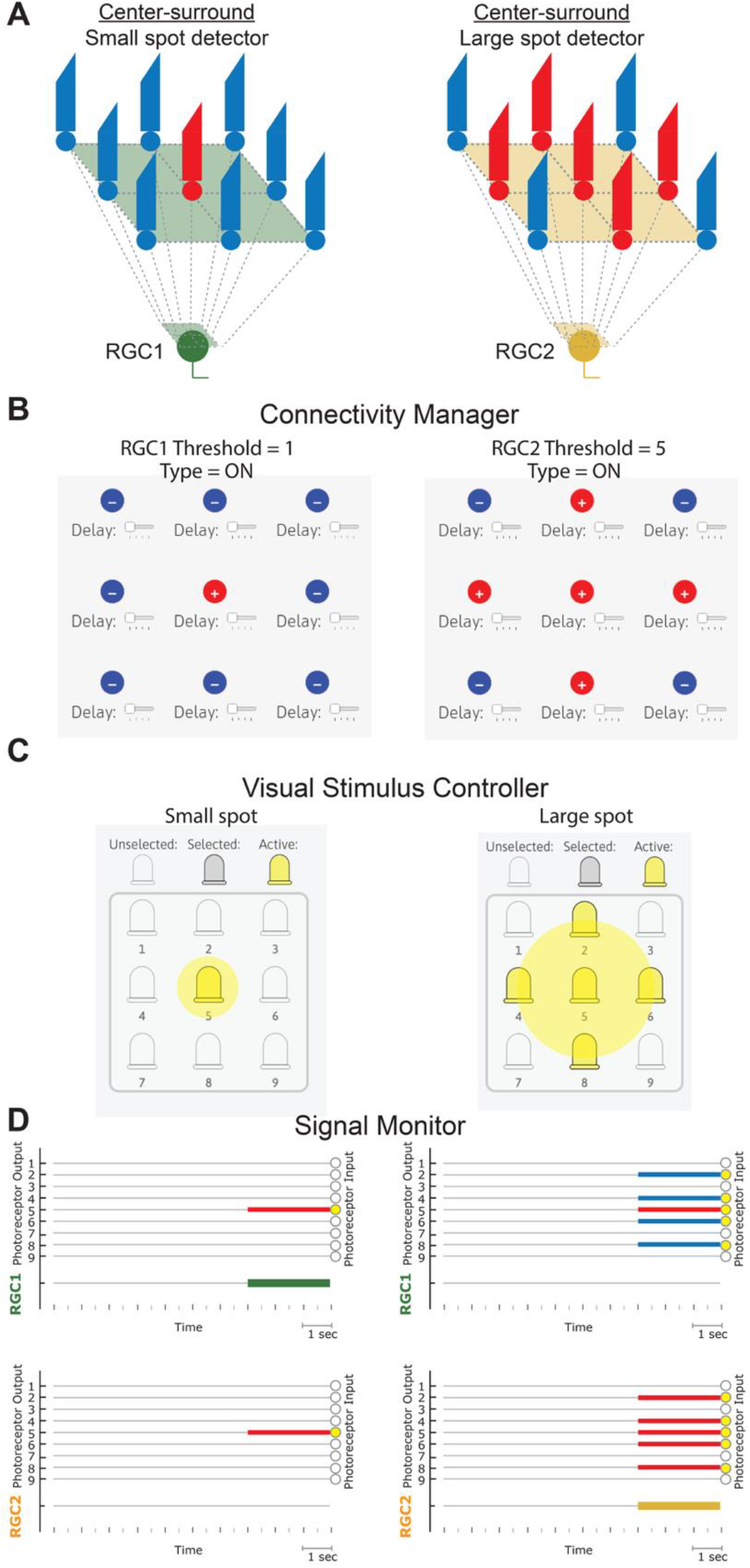
Lesson 1 – Center-surround receptive fields. **A**, Example circuits for ON center-surround RFs with preferences for small (*left*) and large (*right*) spots. Example settings for the Connectivity Manager (**B**) and the Visual Stimulus Controller (**C**). **D**, Example Signal Monitor output for the settings outlined above, with the stimulus LEDs turned on for 3 s (small spot, *left*; large spot, *right*). representing a letter from the alphabet (**Figure 4**). Users are also provided with a cipher containing information they need to use to solve the problem (**Figure 4**). The cipher indicates what the feature preference should be for the two model ON RGCs and then indicates which letters of the alphabet correspond to neither ganglion cell being activated, one or the other ganglion cell being activated in isolation, or both ganglion cells being activated together. Once the Connectivity Manager is set according to the cipher (and in a manner that allows for the possibility that both RGCs can be co-activated), users present RetINaBox with the indicated visual stimuli using the Visual Stimulus Tool or shapes cut out of paper/cardboard. Based on the output of the ganglion cells to each visual stimulus, users enter the corresponding letters into the GUI to solve the code (**Figure 4**). RetINaBox is packaged with multiple codes that users can try to crack.

**Figure 4.**
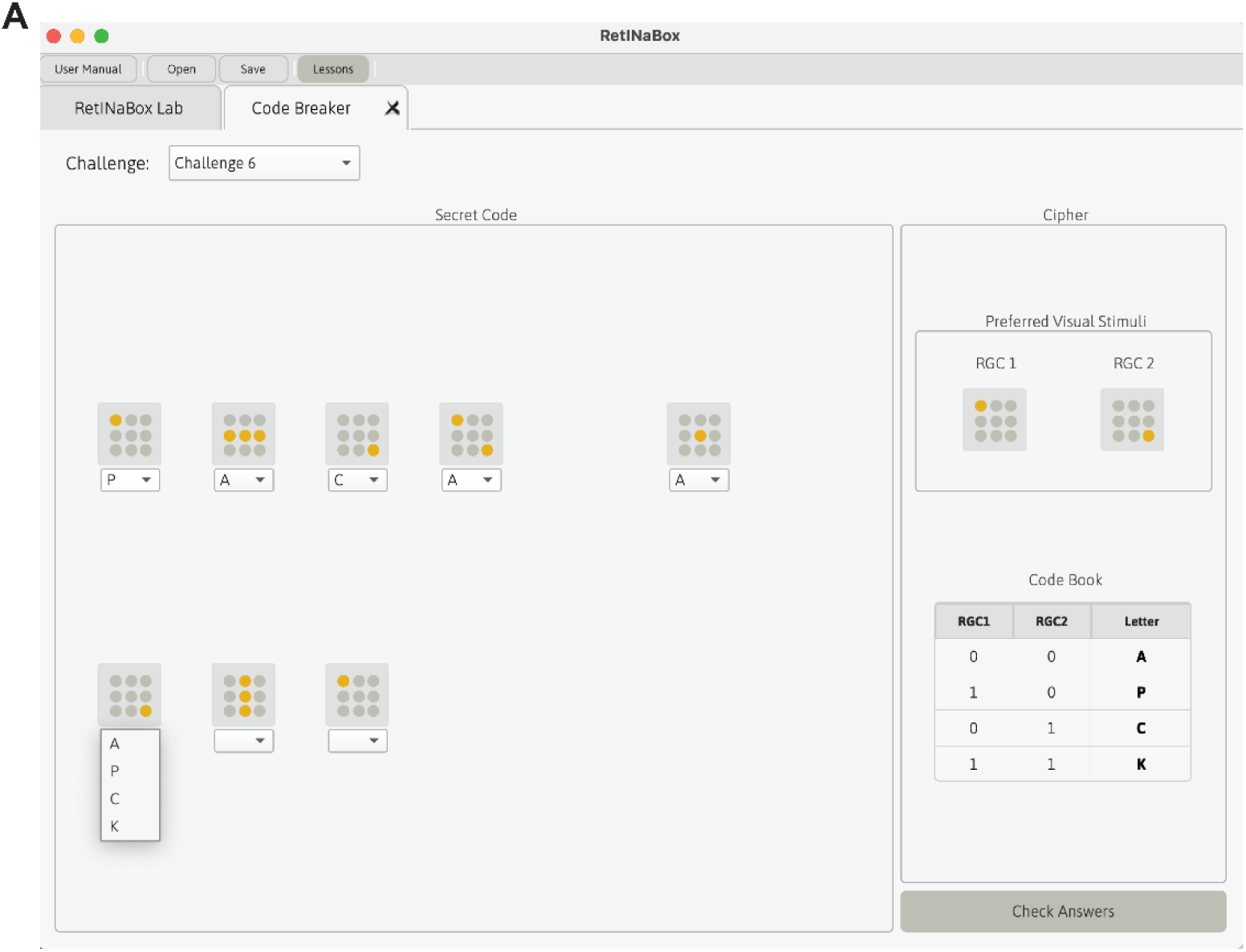
Lesson 1 - Code breaker activity with center-surround receptive fields. **A**, A visualization of the code breaker GUI tab related to Lesson 1. Users must use the cipher (*right*) to correctly set up the Connectivity Manager for the two RGCs to obtain the indicated feature selective responses. Next, users place the visual stimuli indicated on the left into RetINaBox and use the cipher instructions (*bottom right*) to transcribe the RetINaBox output into the correct letter for each visual stimulus in the code.

Lesson 1 ends with users solving a code breaking game, which gets them to apply concepts related to center-surround receptive fields to solve the problem. Upon selecting a code breaking challenge in the GUI, users are provided with a code in the form of a series of visual stimuli, with each visual stimulus

#### Lesson 2 – Build a shape detector with orientation selective receptive fields

Despite receiving its main sensory input from retinal ganglion cells^9,10^—via LGN relay neurons that also tend to possess center-surround receptive fields^26,28^— recordings from primary visual cortex (V1) showed that most V1 neurons do not possess center-surround receptive fields. Instead, many V1 neurons exhibit orientation selective tuning^11,30,31^, meaning that they are optimally activated by an extended edge (or line) in a specific part of the visual field, aligned in a specific orientation. It was posited that such orientation selective responses arise when a single V1 neuron pools inputs from multiple center-surround LGN neurons whose receptive fields are spatially offset along a line^11,29^. While early work in cats and primates found that orientation selectivity appears first in V1^11,30,31^ (but see also^32^), subsequent work in other species, including rabbits^33^ and mice^34,35^, has found that orientation selectivity can also arise at the level of retinal ganglion cells. Such orientation selective receptive fields are thought to be an efficient way to sample visual statistics in the natural world^36,37^. Orientation selective signals can then be combined in downstream neurons in various ways, resulting in spatially invariant orientation selective cells (if a downstream neuron receives input from multiple orientation selective cells with the same orientation preference but with spatially offset receptive fields^11^) and in cells that respond to combinations of edges/lines, which can in turn serve as building blocks for visual object detectors^38^.

Lesson 2 of RetINaBox begins by asking users to build a retinal ganglion cell with an orientation selective receptive field organization. The first goal is to connect the photoreceptors to RGC1 in such a way that it will respond when a specific set of 3 adjacent photoreceptors are activated (i.e. a 3-LED-long line) but not when any other arrangements of photoreceptors are activated (**Figure 5A,B**). Then, users are asked to generate a second orientation selective cell with a preference for the same line as RGC1, but with a preference for a dark line (i.e. build 2 RGCs with the same orientation preference but one with an ON preference and the other with an OFF preference). Next, the user is tasked with setting up RGC2 to exhibit a preference for a line of a different orientation than RGC1 (**Figure 5A,B**). By generating 2 RGCs with different orientation selective properties, RetINaBox helps users understand that each photoreceptor can contribute to distinct parts of each RGC’s receptive field. The next goal is to generate two RGCs with preferences for lines of the same orientation, but of different thicknesses, which demonstrates differences in spatial frequency tuning. Then, users are tasked with generating 2 orientation selective cells with preferences for a bright line of the same orientation and same spatial location, but of different lengths. This helps users learn about the concept of end-stopping^39^, which is thought to be important for enabling encoding of high curvature features in visual scenes^40^. If users require help solving these problems, they can load preset Connectivity Manager solutions from the “Lessons” tab in the menu bar or consult the lesson plan document.

**Figure 5.**
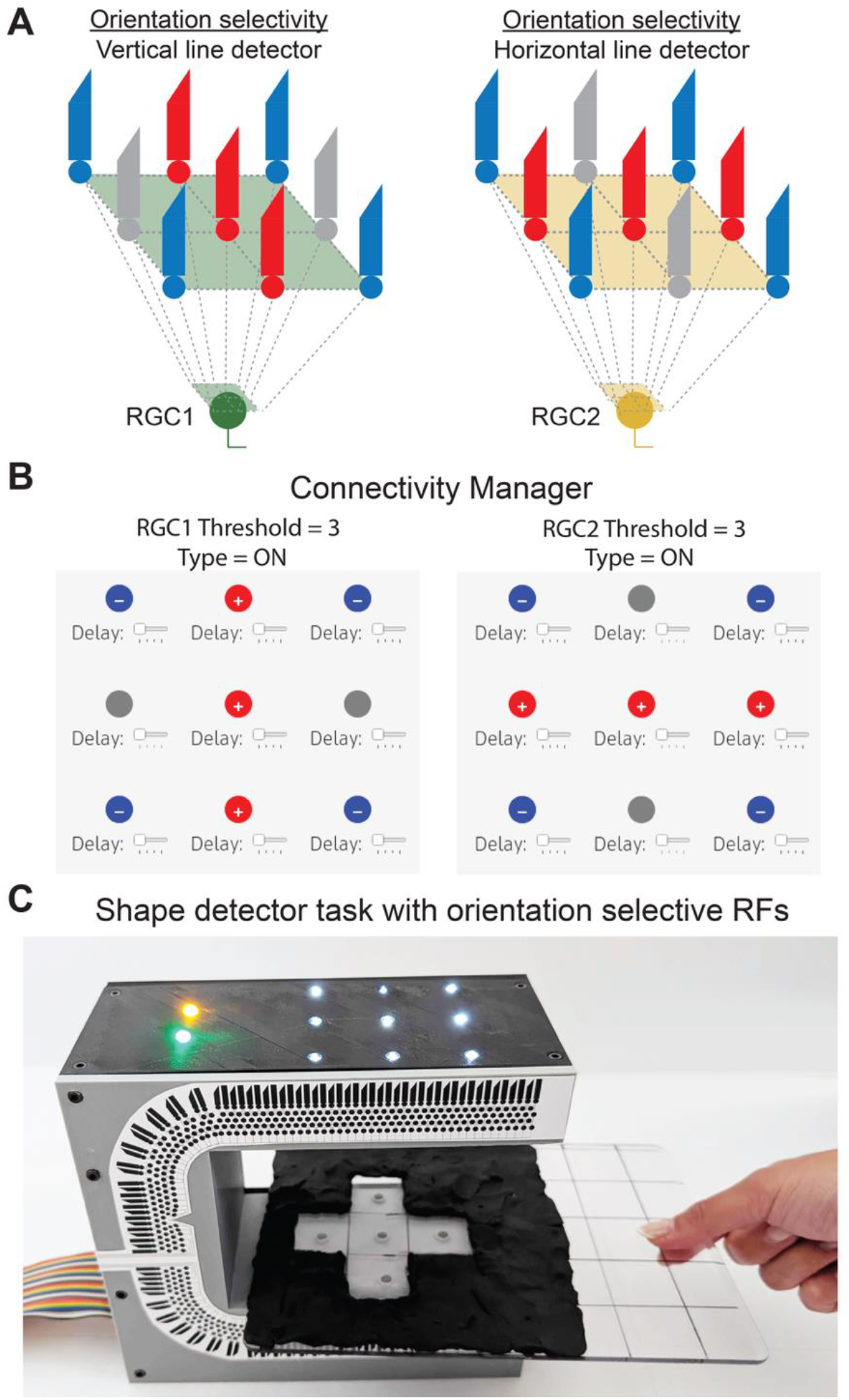
Lesson 2 – Shape detector activity with orientation selective receptive fields. **A**, Example circuits for generating RGCs with orientation selective receptive fields, but preferring lines of different orientations. **B**, Example settings for the Connectivity Manager. **C**, Demonstration of the shape detector activity: the settings in (**B**) were entered into the Connectivity Manager. A shape corresponding to that created by joining the preferred stimuli of RGC1 and RGC2 was made with the Visual Stimulus Tool, which was then placed in the correct spatial location between RetINaBox’s stimulation LEDs and photoreceptor array. Both RGCs are activated at the same time: both yellow and green LEDs are activated.

Lesson 2 ends with users being asked to build a shape detector, which gets users to apply concepts related to orientation selective receptive fields to solve the problem. First, users need to optimize two ON ganglion cells to be selective for lines of different orientations at specific locations over the photoreceptor array, with the combination of the two lines representing a shape (**Figure 5C,D**). Then, users are instructed in basic electronics (see the user manual) to connect the digital outputs of the two ganglion cells (3.3V outputs located on the rear of RetINaBox) to generate a buzzer that is activated via an AND gate, meaning that both ganglion cells need to be activated for the buzzer to sound. This is meant to be analogous to the way that some neuroscientists work, when they listen to their experiments in real-time by playing their electrophysiological recordings through a speaker (which can be helpful for quickly finding stimuli that activate neurons).

#### Lesson 3 – Play a video game with direction selective receptive fields

David Hubel and Torsten Wiesel discovered that in V1 of cats, some cells, aside from being orientation selective, were also direction selective^11^. Soon afterward, direction selective ganglion cells were described in the rabbit retina^12,13^. Direction selective retinal ganglion cells were subsequently found in a variety of species, including mice^41,42^ and primates^43,44^. Direction selective visual responses have been particularly well studied in the fly retina^45^. Direction selective responses can help to stabilize eye/head movements with respect to the visual scene^46–48^, and help an animal distinguish between external and self-generated movements in the visual scene^49,50^. Various models have been proposed and discovered in the brain for generating direction selective responses, including spatially offset excitation/inhibition and spatially asymmetric delays in inputs along the preferred-null axis^15^.

Lesson 3 of RetINaBox asks users to build a retinal ganglion with a direction selective receptive field organization. The goal is to connect the photoreceptors to a retinal ganglion cell in such a way that it will respond when the user moves a visual stimulus (e.g. the user’s hand) in one direction over the photoreceptor array, but will not respond when the same visual stimulus is moved in the opposite direction (**Figure 6A,B**). The next goal is to generate a second ganglion cell with a preference for motion in the opposite direction (**Figure 6A,B**). By generating 2 RGCs with different direction selective properties, users learn that individual photoreceptors can contribute to distinct parts of each RGC’s receptive field. Next, users are instructed to generate 2 ganglion cells, each with a preference for the same visual stimulus moving in the same direction, but at different speeds. This exposes users to the concept of temporal frequency tuning for motion. If users need assistance solving these problems, they can refer to example solutions by loading presets from the “Lessons” tab in the menu bar.

**Figure 6.**
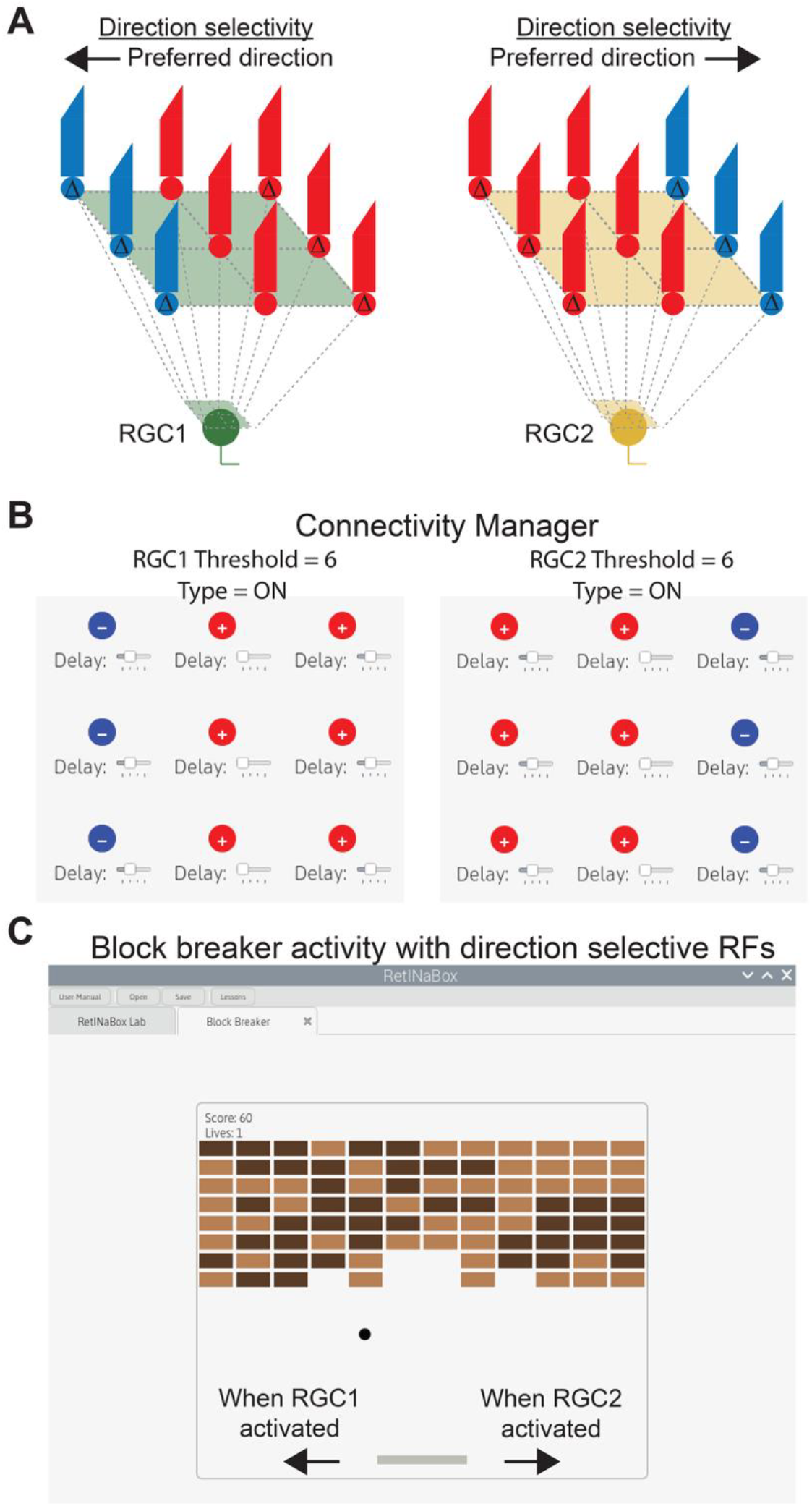
Lesson 3 – Block breaker video game with direction selective receptive fields. **A**, Example circuits for generating direction selective RGCs with preferred directions for leftward (*left*) and rightward (*right*) motion. **B**, Example settings for the Connectivity Manager. **C**, The RetINaBox GUI can be used to load a block breaker video game that takes its left and right input commands from the activity of RGC1 and RGC2, respectively.

Lesson 3 ends with users being asked to build a virtual brain machine interface, which tasks users with generating robust direction selective detectors and applying the output of their retina to control a video game, via left and right commands. First, users need to optimize the two ganglion cells to be selective for motion in opposite directions, as outlined above (**Figure 6A,B**). Then, users are instructed to open a pre-packaged video game in the GUI (**Figure 6C**) which uses the ganglion cell outputs as the digital inputs to the game.

#### Lesson 4 – Discovery mode: What does RetINaBox want to see?

Lessons 1-3 centered around well-established feature selective computations in the visual system. These feature selective properties were discovered by scientists presenting varied sets of visual stimuli while recording from neurons, until eventually the stimulus that best activated each neuron was found. Once the preferred visual stimulus of a neuron was found, scientists then turned their attention to discovering the circuit wiring that enabled such a feature selectivity. To replicate this discovery aspect of scientific exploration, Lesson 4 tasks users with discovering the visual features that drive mystery ganglion cells in RetINaBox and uncovering the circuits that underlie their feature selective responses (**Figure 7A-C**).

**Figure 7.**
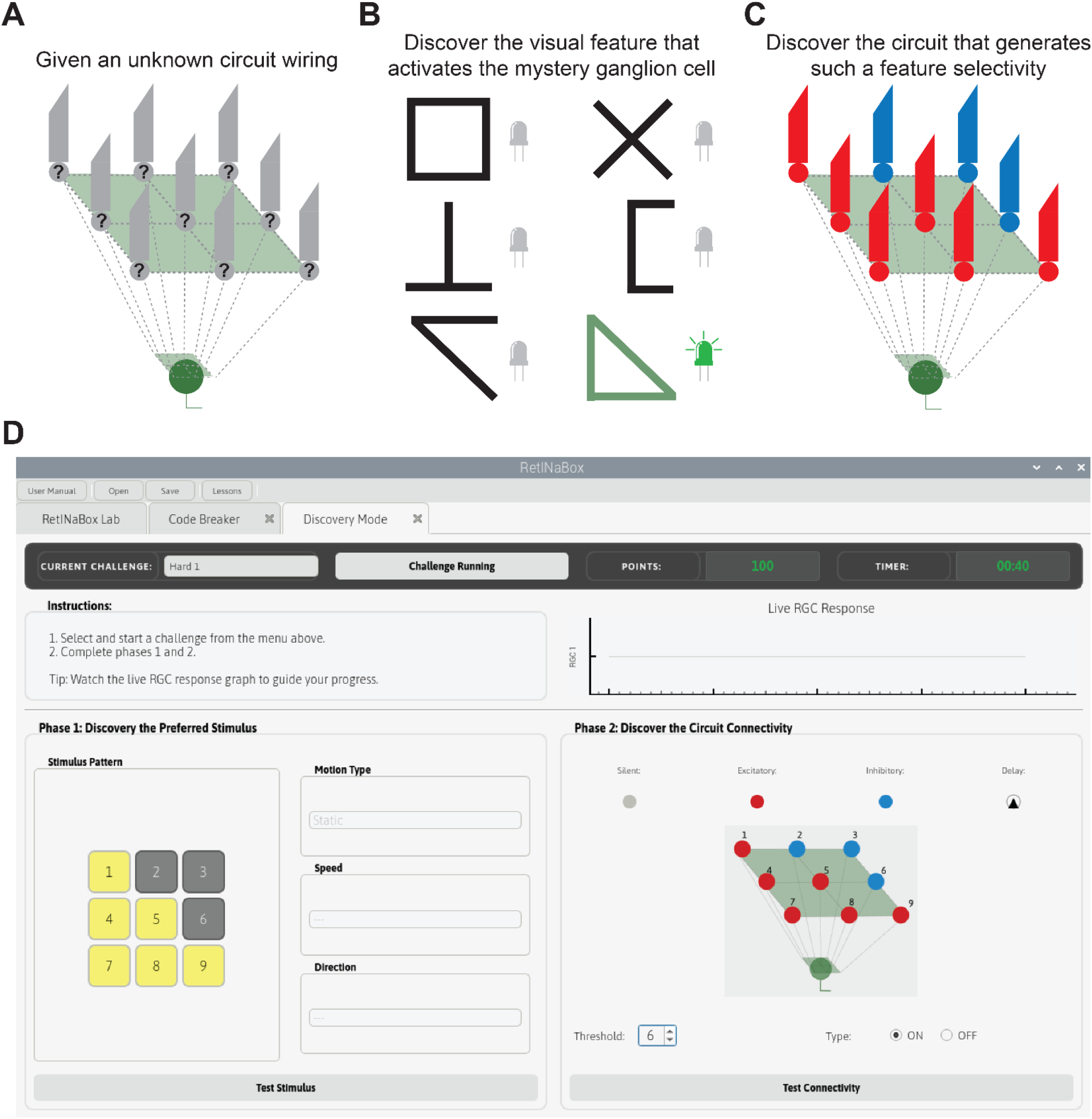
Lesson 4 – Discovery mode. Users select a mystery ganglion cell (**A**), test various visual stimuli to discover the preferred stimulus of the mystery ganglion cell (**B**), and discover the circuit connectivity that generates such a feature selectivity (**C**), all using the Discovery Mode GUI tab (**D**).

Discovery Mode opens a new GUI tab (**Figure 7D**). Users start by selecting a mystery ganglion cell from a drop-down list. Mystery ganglion cells in the Discovery Mode menu section are organized based on difficulty (easy, medium, and hard). In Phase 1, users are tasked with figuring out which visual stimulus activates the mystery ganglion cell—by using the Visual Stimulus Tool, or their hands, or patterns cut out of paper/cardboard—to test various stimuli at different positions, orientations, and directions/speeds of movement over RetINaBox’s photoreceptor array. Upon identifying the preferred visual stimulus for a mystery ganglion cell, users can verify in the software that they have correctly identified the target visual stimulus (**Figure 7D**). Once users have found the target stimulus, in Phase 2 users must discover the connectivity settings that enable a ganglion cell to be selective for the target visual stimulus they just discovered (**Figure 7D**).

## Discussion

Taking inspiration from groups that sought to develop interactive learning tools targeted towards neuroscience education^51,52^, and those who sought to make electronic models of the visual system^53,54^, we developed RetINaBox. However, unlike these previous systems, which either allowed users to interact with model neurons with biophysically realistic properties^51^, or enabled users to provide visual stimuli to a complex electrical device that responded in a similar manner as visual neurons in the brain^53,54^, here we specifically tailored RetINaBox to let users learn about several concepts in visual neuroscience—ON/OFF, center-surround, orientation selective and direction selective feature selective tuning properties—through the act of exploration and discovery to try to recreate, in a classroom setting, the lab experience of recording from visual neurons.

RetINaBox is an electronic visual stimulation/detection device paired with a computer. RetINaBox comes with 4 detailed lesson plans. In each lesson users are encouraged to complete the tasks by reflecting on which visual feature detectors they are trying to build and which circuit components are available to them. Users are then guided to develop and test hypotheses while working towards a circuit model for each distinct feature detector. While going through the lessons and building/testing circuits, users learn important concepts in neuroscience, including excitatory and inhibitory synaptic connectivity, response thresholds, spatiotemporal processing, and parallel processing.

As outlined above in the section detailing the design of RetINaBox, it is important to note that RetINaBox represents a simplification of actual biological circuits (e.g. we bypass bipolar/horizontal/amacrine cells in the retina and replace them with sign, delay and ON/OFF functions). Nonetheless, we believe that RetINaBox provides a useful heuristic for approximating visual neuroscience experiments for educational purposes insomuch as it allows users to focus on general concepts related to feature selective tuning preferences. These concepts can then be applied towards understanding how such feature selective computations are implemented in various ways in different parts of the visual system. Importantly, to ensure that RetINaBox users are not left with an incorrect understanding of how the actual visual system works, the lesson plans include details about the biological circuits that underlie the various visual feature detectors covered in lessons 1-3.

Regarding its usage, we designed RetINaBox for neuroscience outreach events, whether it be with students (high school or undergrad) or the general public. Regarding graduate students, while someone who is specifically studying circuit processing in the retina or visual cortex may find RetINaBox to be overly simplistic, we believe it can be a useful tool for conveying concepts in visual processing to graduate students working in molecular, genetic, and clinical aspects of vision, who often do not have backgrounds in circuits and systems neuroscience. Thus, we believe that RetINaBox can be a useful learning tool for various types of users. To aid in its use, in addition to lesson plans we also provide teaching slides for RetINaBox, meaning that RetINaBox can be easily incorporated into existing neuroscience classes or quickly set up as an outreach activity. It should be noted that we did not design RetINaBox to replace other pedagogical tools (such as lectures, or videos, or quizzes), but as an additional tool to complement the existing repertoire of neuroscience related teaching methods.

Lastly, to ensure that RetINaBox is accessible to as many people as possible, we built it with simple, low-cost parts and open-source software. All custom-written code, as well as detailed instructions for building and running the system, are available in the user manual. Seeing as RetINaBox is designed from simple electronic components (photodiodes and LEDs) and a few 3D printed parts, it can be further expanded or altered by RetINaBox users for their own specific use cases. Along these lines, while RetINaBox uses most of the Raspberry Pi’s GPIO ports, several remain unused and could be leveraged for additional purposes as users see fit. Furthermore, while we tried to keep RetINaBox as simple as possible while also providing a versatile system that could be used to explore the topics of ON/OFF, center-surround, orientation selective and direction selective processing, the GUI software is open-source and thus can be edited as users see fit if they wish to add new features. As one example, if a user wanted to include additional details about temporal processing, they could add time constant variables to enable responses to be either sustained or transient. As another example, if a user wanted to add more ganglion cells to better highlight parallel processing and population codes, this would be possible. As such, while completely functional as is, RetINaBox could be expanded upon in various ways as users see fit.

## Contributions

The software was developed by B.B. and S.T. and all code was written by B.B. RetINaBox’s electronic retina was designed by B.B., F.A.A. and S.T. The 3D printed components were designed by F.A.A. and S.T. RetINaBox was assembled by F.A.A. The lesson plans were developed by E.C. and S.T and made by E.C. The user manual was written by B.B., F.A.A., and S.T. Beta testing was performed by V. B., M. Liu, and M. Loukine. Translations of the lesson plans and user manual were undertaken by F.A.A., V.B., M. Loukine, M.W., and A.V. Teaching slides were generated by E.C. The RetINaBox logo was designed by B.B. and the decorative retina image on our RetINaBox (as seen in Figure 5C) was designed by V.B. B.R. assisted with conceptual development of the project. The manuscript was written by S.T.

## Acknowledgements

We thank A. Krishnaswamy for critical discussions. We thank H. Velde for noting that “retina in a box” featured “ina in a,” which prompted us to shorten the name from Retina-In-A-Box to RetINaBox. We thank the following funding sources: a Vision Sciences Research Network Merit Scholarship to F.A.A.; a National Sciences and Engineering Research Council (NSERC) and Fonds de Recherche du Québec (FRQ) scholarship to E.C.; a Canadian Institutes of Health Research (CIHR) PhD fellowship to M. Liu; an NSERC Undergraduate Summer Research Award, a Tanenbaum Open Science Institute Launchpad award, and an NSERC graduate student scholarship M. Loukine; a Tomlinson PhD fellowship to M.W.; CIFAR Canada AI Chair (Learning in Machine and Brains Fellowship) to B.R.; a Canada Research Chair, an Alfred P. Sloan Foundation Research Fellowship, and NSERC Discovery Grants (RGPIN-2018-03852 and RGPIN-2025-05567) to S.T. We also acknowledge outreach funding from Brain Canada, the Vision Sciences Research Network, and the Canadian Association for Neuroscience.

Custom code, 3D-print files, the user manual, and the lesson plans can be found here: https://github.com/Trenholm-Lab/RetINaBox

## References

1. Lettvin, J. Y., Maturana, H. R., McCulloch, W. S. & Pitts, W. H. What the Frog’s Eye Tells the Frog’s Brain. Proc. IRE 47, 1940–1951 (1959).

2. The Man Who Tried to Redeem the World with Logic - Nautilus. https://nautil.us/the-man-who-tried-to-redeem-the-world-with-logic-235253/.

3. Hubel, D. H. & Wiesel, T. N. Receptive fields of single neurones in the cat’s striate cortex. J. Physiol. 148, 574–591 (1959).

4. Hubel, D. H. & Wiesel, T. N. Chapter 7 - Our first paper, on cat cortex, 1959. in Brain and Visual Perception 59–67 (Oxford University Press).

5. Trenholm, S. & Krishnaswamy, A. An Annotated Journey through Modern Visual Neuroscience. J. Neurosci. Off. J. Soc. Neurosci. 40, 44–53 (2020).

6. Hartline, H. K. THE RESPONSE OF SINGLE OPTIC NERVE FIBERS OF THE VERTEBRATE EYE TO ILLUMINATION OF THE RETINA. Am. J. Physiol.-Leg. Content 121, 400–415 (1938).

7. Famiglietti, E. V. & Kolb, H. Structural Basis for ON-and OFF-Center Responses in Retinal Ganglion Cells. Science 194, 193–195 (1976).

8. Slaughter, M. M. & Miller, R. F. 2-amino-4-phosphonobutyric acid: a new pharmacological tool for retina research. Science 211, 182–185 (1981).

9. Barlow, H. B. Summation and inhibition in the frog’s retina. J. Physiol. 119, 69–88 (1953).

10. Kuffler, S. W. DISCHARGE PATTERNS AND FUNCTIONAL ORGANIZATION OF MAMMALIAN RETINA. J. Neurophysiol. 16, 37–68 (1953).

11. Hubel, D. H. & Wiesel, T. N. Receptive fields, binocular interaction and functional architecture in the cat’s visual cortex. J. Physiol. 160, 106–154 (1962).

12. Barlow, H. B., Hill, R. M. & Levick, W. R. RETINAL GANGLION CELLS RESPONDING SELECTIVELY TO DIRECTION AND SPEED OF IMAGE MOTION IN THE RABBIT. J. Physiol. 173, 377–407 (1964).

13. Barlow, H. B. & Levick, W. R. The mechanism of directionally selective units in rabbit’s retina. J. Physiol. 178, 477–504 (1965).

14. Antinucci, P. & Hindges, R. Orientation-Selective Retinal Circuits in Vertebrates. Front. Neural Circuits 12, (2018).

15. Mauss, A. S., Vlasits, A., Borst, A. & Feller, M. Visual Circuits for Direction Selectivity. Annu. Rev. Neurosci. 40, 211–230 (2017).

16. Krishnaswamy, A. & Trenholm, S. The Retina. in The Open Brain (2023).

17. Ahnelt, P. K. The photoreceptor mosaic. Eye 12, 531–540 (1998).

18. Ahnelt, P. K. & Kolb, H. The mammalian photoreceptor mosaic-adaptive design. Prog. Retin. Eye Res. 19, 711–777 (2000).

19. Adrian, E. D. & Matthews, R. The action of light on the eye. J. Physiol. 63, 378–414 (1927).

20. Bipolar Cell Pathways in the Vertebrate Retina by Ralph Nelson and Victoria Connaughton – Webvision. https://www.webvision.pitt.edu/book/part-v-phototransduction-in-rods-and-cones/bipolar-cell-pathways-in-the-vertebrate-retina/.

21. Schiller, P. H., Sandell, J. H. & Maunsell, J. H. Functions of the ON and OFF channels of the visual system. Nature 322, 824–825 (1986).

22. Baylor, D. A., Fuortes, M. G. & O’Bryan, P. M. Receptive fields of cones in the retina of the turtle. J. Physiol. 214, 265–294 (1971).

23. Kawai, F. Certain retinal horizontal cells have a center-surround antagonistic organization. J. Neurophysiol. 128, 1337–1343 (2022).

24. Werblin, F. S. & Dowling, J. E. Organization of the retina of the mudpuppy, Necturus maculosus. II. Intracellular recording. J. Neurophysiol. 32, 339–355 (1969).

25. Nelson, R. AII amacrine cells quicken time course of rod signals in the cat retina. J. Neurophysiol. 47, 928–947 (1982).

26. Singer, W. & Creutzfeldt, O. D. Reciprocal lateral inhibition of on- and off-center neurones in the lateral geniculate body of the cat. Exp. Brain Res. 10, 311–330 (1970).

27. Molnar, A., Hsueh, H.-A., Roska, B. & Werblin, F. S. Crossover inhibition in the retina: circuitry that compensates for nonlinear rectifying synaptic transmission. J. Comput. Neurosci. 27, 569– 590 (2009).

28. Hubel, D. H. & Wiesel, T. N. Integrative action in the cat’s lateral geniculate body. J. Physiol. 155, 385-398.1 (1961).

29. Angelucci, A. & Trenholm, S. Primary Visual Cortex. in Adler’s Physiology of the Eye 612–626 (Elsevier).

30. Hubel, D. H. & Wiesel, T. N. Receptive fields and functional architecture of monkey striate cortex. J. Physiol. 195, 215–243 (1968).

31. Hubel, D. H. & Wiesel, T. N. Ferrier lecture. Functional architecture of macaque monkey visual cortex. Proc. R. Soc. Lond. B Biol. Sci. 198, 1–59 (1977).

32. Passaglia, C. L., Troy, J. B., Rüttiger, L. & Lee, B. B. Orientation sensitivity of ganglion cells in primate retina. Vision Res. 42, 683–694 (2002).

33. Levick, W. R. Receptive fields and trigger features of ganglion cells in the visual streak of the rabbit’s retina. J. Physiol. 188, 285–307 (1967).

34. Baden, T. et al. The functional diversity of retinal ganglion cells in the mouse. Nature 529, 345– 350 (2016).

35. Nath, A. & Schwartz, G. W. Cardinal Orientation Selectivity Is Represented by Two Distinct Ganglion Cell Types in Mouse Retina. J. Neurosci. 36, 3208–3221 (2016).

36. Olshausen, B. A. & Field, D. J. Emergence of simple-cell receptive field properties by learning a sparse code for natural images. Nature 381, 607–609 (1996).

37. Coppola, D. M., Purves, H. R., McCoy, A. N. & Purves, D. The distribution of oriented contours in the real world. Proc. Natl. Acad. Sci. 95, 4002–4006 (1998).

38. Hesse, J. K. & Tsao, D. Y. The macaque face patch system: a turtle’s underbelly for the brain. Nat. Rev. Neurosci. 21, 695–716 (2020).

39. Hubel, D. H. & Wiesel, T. N. Receptive fields and functional architecture in two nonstriate visual areas (18 and 19) of the cat. J. Neurophysiol. 28, 229–289 (1965).

40. Ponce, C. R., Hartmann, T. S. & Livingstone, M. S. End-Stopping Predicts Curvature Tuning along the Ventral Stream. J. Neurosci. 37, 648–659 (2017).

41. Weng, S., Sun, W. & He, S. Identification of ON–OFF direction-selective ganglion cells in the mouse retina. J. Physiol. 562, 915–923 (2005).

42. Sun, W., Deng, Q., Levick, W. R. & He, S. ON direction-selective ganglion cells in the mouse retina. J. Physiol. 576, 197–202 (2006).

43. Kim, Y. J. et al. Origins of direction selectivity in the primate retina. Nat. Commun. 13, 2862 (2022).

44. Wang, A. Y. M. et al. An ON-type direction-selective ganglion cell in primate retina. Nature 623, 381–386 (2023).

45. Borst, A., Haag, J. & Mauss, A. S. How fly neurons compute the direction of visual motion. J. Comp. Physiol. A 206, 109–124 (2020).

46. Oyster, C. W., Takahashi, E. & Collewijn, H. Direction-selective retinal ganglion cells and control of optokinetic nystagmus in the rabbit. Vision Res. 12, 183–193 (1972).

47. Yoshida, K. et al. A Key Role of Starburst Amacrine Cells in Originating Retinal Directional Selectivity and Optokinetic Eye Movement. Neuron 30, 771–780 (2001).

48. Yonehara, K. et al. Congenital Nystagmus Gene FRMD7 Is Necessary for Establishing a Neuronal Circuit Asymmetry for Direction Selectivity. Neuron 89, 177–193 (2016).

49. Britten, K. H. Mechanisms of Self-Motion Perception. Annu. Rev. Neurosci. 31, 389–410 (2008).

50. Sabbah, S. et al. A retinal code for motion along the gravitational and body axes. Nature 546, 492– 497 (2017).

51. Dragly, S.-A. et al. Neuronify: An Educational Simulator for Neural Circuits. eNeuro 4, (2017).

52. Book: How Your Brain Works. Backyard Brains https://backyardbrains.com/products/book-how-your-brain-works.

53. Delbrück, T. & Liu, S.-C. A silicon early visual system as a model animal. Vision Res. 44, 2083– 2089 (2004).

54. Li, G., Talebi, V., Yoonessi, A. & Baker, C. L. A FPGA real-time model of single and multiple visual cortex neurons. J. Neurosci. Methods 193, 62–66 (2010).

